# Regulators H-NS and LeuO inversely control swarming motility and biofilm formation in *Vibrio parahaemolyticus*

**DOI:** 10.1101/2024.01.09.574875

**Authors:** S.M. Bhide, J.G. Tague, J. A. Andrews, K.E. Boas Lichty, E.F. Boyd

## Abstract

*Vibrio parahaemolyticus* is a halophile present in marine environments worldwide and is a leading cause of bacterial seafood-borne gastroenteritis. Free living *Vibrio parahaemolyticus* planktonic cells can either attach to surfaces to form swarming cells or develop into a sessile three-dimensional biofilm structure. Swarming motility requires lateral flagella (*laf* operon) and the expression of the surface sensing operon *scrABC* to produce a spreading cauliflower colony morphology. Biofilms are formed from capsule polysaccharide (CPS) encoded by the *cpsA-K* operon that is positively regulated by CpsR and CpsQ. In enteric bacteria, H-NS is a global gene silencer and LeuO is an antagonist of H-NS. In this work, we examined the role of these regulators in the decision between swarming and biofilm behaviors using deletion mutants of *leuO, hns,* and a double deletion *leuO/hns*. The wild type and Δ*leuO* strains produced swarming colonies whereas Δ*hns* produced a hyper swarming whereas in Δ*leuO/*Δ*hns,* the phenotype reverted to wild type. Transcriptional reporter assays of P*lafB-gfp* and P*scrA*-*gfp* showed significantly increased fluorescence in Δ*hns* compared to wild type. In the Δ*leuO/*Δ*hns* mutant, P*lafB-gfp* fluorescence reverted to wild type levels and P*scrA-gfp* showed increased fluorescence compared to wild type. In CPS assays, Δ*leuO* had a less dense rugose morphology compared to wild type and Δ*hns* produced a smooth colony, which also produced significantly less biofilm. Δ*leuO/*Δ*hns* had an opaque morphology and produced significantly more biofilm. Reporter expression assays of P*cpsA-gfp* and P*cpsR-gfp* confirmed the roles of both H-NS and LeuO in CPS and biofilm formation.

**Importance:** This study determined the role of LeuO and H-NS in controlling the decision between two surface based behaviors, swarming motility and sessile biofilm formation in *V. parahaemolyticus*. The effects of deletions of *leuO*, *hns,* and a double Δ*leuO/*Δ*hns* mutant showed that H-NS was a negative regulator of swarming, but a positive regulator of biofilm formation. The mechanism of this control was in part due to H-NS inhibition of LeuO at loci required for swarming and biofilm formation.

## INTRODUCTION

*Vibrio parahaemolyticus* is an abundant marine heterotrophic bacterium that is a significant pathogen of humans and marine fauna that has shown extensive geographic spread to previously unoccupied waters (1–4). *Vibrio parahaemolyticus* can live as a planktonic swimming cell or attach to biotic and abiotic surfaces either as a part of motile swarmer colony or as a sessile biofilm (5). *Vibrio parahaemolyticus* produces two flagellum systems, a single polar flagellum in liquid media for swimming motility and numerous lateral flagella on surfaces for swarming motility producing forming spreading cauliflower shaped colonies (5–11). In *V. parahaemolyticus*, the lateral flagella are encoded by the *laf* operon and LafK is the major positive regulator of *laf* gene expression and is an activator of the σ-factor RpoN (8, 10, 11). The surface sensing operon *scrABC*, is also required for swarming motility, and encodes a phosphodiesterase that degrades intracellular bis-(3′-5′)-cyclic dimeric guanosine monophosphate (c-di-GMP) (12–16). Swarming motility does not require c-di-GMP, but deletion of *scrABC* and *scrG*, which encodes an additional phosphodiesterase, significantly decreases *laf* gene expression and prevents swarming motility (12–17). In *V. parahaemolyticus* and many other bacterial species, a high concentration of c-di-GMP is required for the switch from motility to sessility (12–23) (24).

A biofilm is three-dimensional surface-based bacterial aggregation and likely a preferred lifestyle for most bacteria, as it protect cells from nutrient limitations, antimicrobials, abiotic stress conditions, host immune responses as well as protozoan grazing (25–27). Biofilms are self-sustaining bacterial communities embedded in a matrix of extracellular polymeric substances (EPS) such as proteins and extracellular capsule polysaccharides (CPS) (25–27). In *V. parahaemolyticus,* CPS is a major component of biofilm and is encoded by the *cpsA-K* operon. The *cps* operon is directly positively regulated by CpsQ, which is positively regulated by CpsR (14, 28–30). CpsQ is a member of the CsgD family of c-di-GMP binding proteins and requires the second messenger c-di-GMP for activation (13, 14).

H-NS (histone-like nucleoid structuring) is a small (135 amino acid) abundant nucleoid-associated protein (NAP) involved in bacterial chromosome organization and in *E. coli,* is characterized as a global gene silencer (31–37). H-NS contains an N-terminal oligomerization domain and a C-terminal DNA binding domain and preferentially binds to AT-rich regions such as intergenic regions and horizontally acquired DNA (31–37). In *E. coli*, H-NS silencing of gene expression has been shown to be inhibited via the anti-silencing activity of proteins such as LeuO (37–39). LeuO anti-silencing activity can be achieved either by blocking H-NS oligomerization sites or by removing H-NS from bound DNA (37–39). H-NS and LeuO are antagonists at many loci in *E. coli, S. enterica,* and *V. cholerae*, regulating a number of phenotypes in these species (37, 38, 40–43).

In *V. parahaemolyticus*, an *hns* deletion mutant was shown to be defective in CPS, biofilm, and swimming motility, but was swarming proficient (29, 44, 45). McCarter and colleagues showed that LeuO (named CalR) was a negative regulator of swarming and *laf* gene expression (46). LeuO in *V. parahaemolyticus* is positively regulated by the global regulator ToxR and is required for acid tolerance (47). Also, a recent study showed an epistatic relationship between LeuO and H-NS in the control of osmotic stress in this species (48).

In the present work, the role of LeuO and H-NS in controlling the decision between swarming and sessile biofilm formation, two mutually exclusive behaviors was determined. The effects of deletions of *leuO* and *hns* in swarming motility, capsule production, and biofilm formation was examined. Epistasis was determined by examining a double Δ*leuO/*Δ*hns* mutant. Overall, we found that the Δ*hns* mutant had a hyper swarming phenotype, but produced significantly less CPS and biofilm compared to wild type. The Δ*leuO* mutant showed less significant changes in swarming behavior and CPS production and no change in biofilm formation. However, the Δ*leuO*/Δ*hns* double mutant demonstrated more clearly the role of LeuO in these phenotypes. In the swarming assay, the double mutant had a phenotype similar to wild type indicating that in the Δ*hns* mutant the hyper swarming phenotype was likely the result of LeuO availability. Reporter expression assays of P*lafB-gfp* and P*scrA-gfp* confirmed H-NS negative role in swarming. In the CPS and biofilm assays, the Δ*hns* mutant produced no CPS and significantly less biofilm formation, Δ*leuO* showed less CPS but no changes in biofilm compared to wild type. The double mutant produced CPS and produced significantly more biofilm indicating H-NS is a positive regulator. P*cpsA-gfp* and P*cpsR-gfp* reporter assays confirmed H-NS role as a positive regulator likely as an inhibitor of *leuO*, a negative regulator of CPS production acting at the *cpsA* and *cpsR* loci.

## RESULTS

### H-NS and LeuO regulate motility in *V. parahaemolyticus*

We wanted to determine the role of LeuO and H-NS in controlling swarming motility and biofilm formation, mutually exclusive behaviors in *V. parahaemolyticus*. To accomplish this, in-frame non-polar deletion of *hns* (VP1133) and *leuO* (VP0350) and a double deletion mutant *leuO/hns* constructed using SOE PCR and allelic exchange were used (48). To ensure that none of the deletions in *V. parahaemolyticus* leads to a growth defect, growth was examined in LB broth with no growth differences observed among strains (data not shown). Swimming motility assays were performed in LB broth 2% NaCl with 0.3% agar incubated at 37°C for 24 h. Measurement of the swimming diameter indicated reduced swimming in Δ*hns* compared to the other strains but this was not statistically significant (data not shown). Next, we performed a swarming motility assay on HI 2% NaCl agar plates incubated at 30°C for 48 h. For wild type, we observed the characteristic “cauliflower-like” colony morphology, Δ*leuO* also showed this morphology with slightly larger colonies (**Fig. 1**). The Δ*hns* strain produced large hyper-swarmer colonies, whereas Δ*leuO*/Δ*hns* showed a colony morphology similar to wild type (**Fig. 1**). These data show an epistatic relationship between LeuO and H-NS with H-NS a negative regulator of swarming.

**Figure 1.**
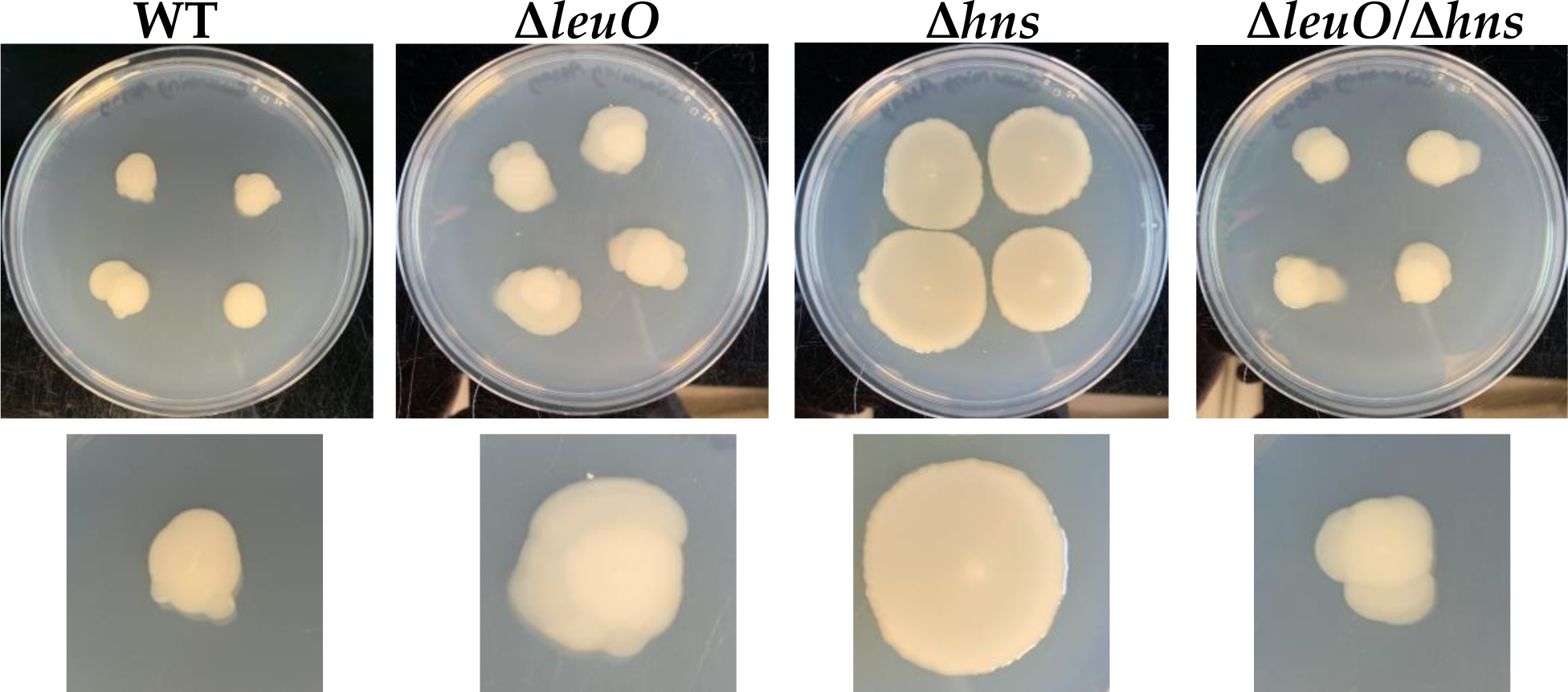
Swarming motility assays. Wild type (WT), Δ*leuO,* Δ*hns,* and Δ*leuO/*Δ*hns* strains inoculated on HI plates supplements 2% NaCl and incubated at 30°C for 48 h. Images are an example of three biological replicates.

In *E. coli*, previous studies showed that H-NS is a negative regulator of *leuO* expression, and LeuO is a negative auto regulator (38, 49). Thus, we were interested to determine whether H-NS is involved in regulating *leuO* in *V. parahaemolyticus* under the conditions examined in this study. To investigate this, we utilized a GFP fluorescence reporter vector pRUP-*gfp*, encoding the *leuO* promoter region P*leuO* upstream of *gfp*. This vector was transformed into wild type, Δ*leuO*, Δ*hns*, and Δ*leuO*/Δ*hns* strains (**Fig. 2**). All strains were grown in HI media with 0.5% NaCl and tetracycline (Tet) at 30°C for 18 h and GFP fluorescence measured. Wild type showed a low level of fluorescence, whereas each of the mutants showed increasing fluorescence levels compared to wild type (**Fig. 2**). The double mutant showed that highest increase indicating that both H-NS and LeuO are negative regulators of *leuO* expression (**Fig. 2**).

**Figure 2.**
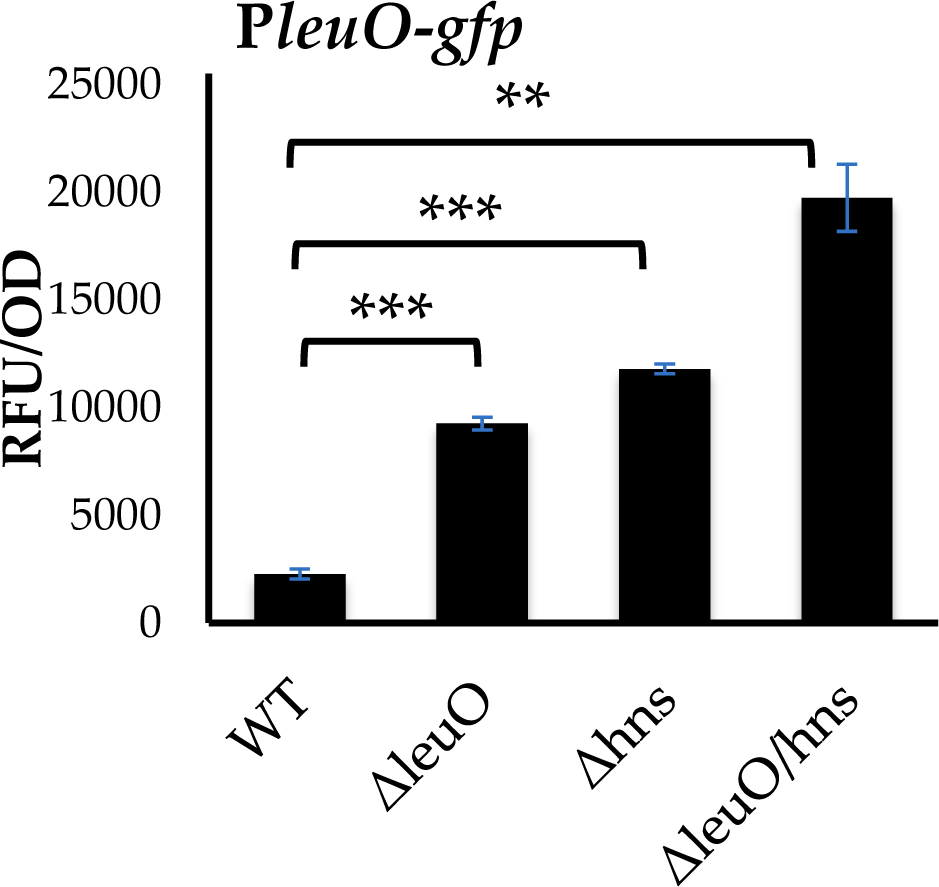
Regulation of *leuO* by LeuO and H-NS. Reporter assays with pRUP*leuO*-*gfp* in wild type (WT), Δ*leuO,* Δ*hns,* and Δ*leuO/*Δ*hns* backgrounds. All strains were grown HI plates supplemented with Tet and 0.5% NaCl at 30°C for 18 h. All assays were performed with at least two biological replicates with three technical replicates. Error bars indicate standard deviation. Unpaired t-test was used to determine the p-value. **p<0.01, ***p<0.001

### H-NS and LeuO counteract each other at the *lafB* and *scrA* loci

Next, we wanted to determine the mechanism of LeuO and H-NS regulation of swarming motility. To accomplish this, we examined the expression of two operons required for the swarming phenotype, the *laf* operon, which encodes the genes for the biosynthesis of lateral flagella, and the *scrABC* operon, which encodes a phosphodiesterase. A GFP-based transcription reporter assay with the promoter region of *lafB* cloned into pRUP to created pRUP*lafB-gfp* and cloned into wild type, Δ*leuO*, Δ*hns* and Δ*leuO*/Δ*hns* mutants was performed (**Fig. 3A**). Transcription was measured by determining GFP fluorescence in cells grown on HI plates with 2% NaCl Tet at 30°C for 18 h. P*lafB*-*gfp* fluorescence in Δ*leuO* was similar to wild type, but Δ*hns* showed increased fluorescence compared to wild-type and Δ*leuO*, but the data was not statistically significant. In the double mutant fluorescence reverted to wild type and Δ*leuO* levels (**Fig. 3A**). The surface-sensing operon, *scrABC,* is one of several loci that control the intracellular concentration of c-di-GMP via the phosphodiesterase activity of ScrC (12, 13, 15). P*scrA-gfp* showed significantly reduced fluorescence in Δ*leuO* compared to wild type whereas Δ*hns* showed significantly increased fluorescence (**Fig. 3B**). The Δ*leuO*/Δ*hns* mutant also showed increased fluorescence compared to Δ*hns*. These data indicate LeuO is epistatic to H-NS, and H-NS negatively impacts swarming motility.

**Figure 3.**
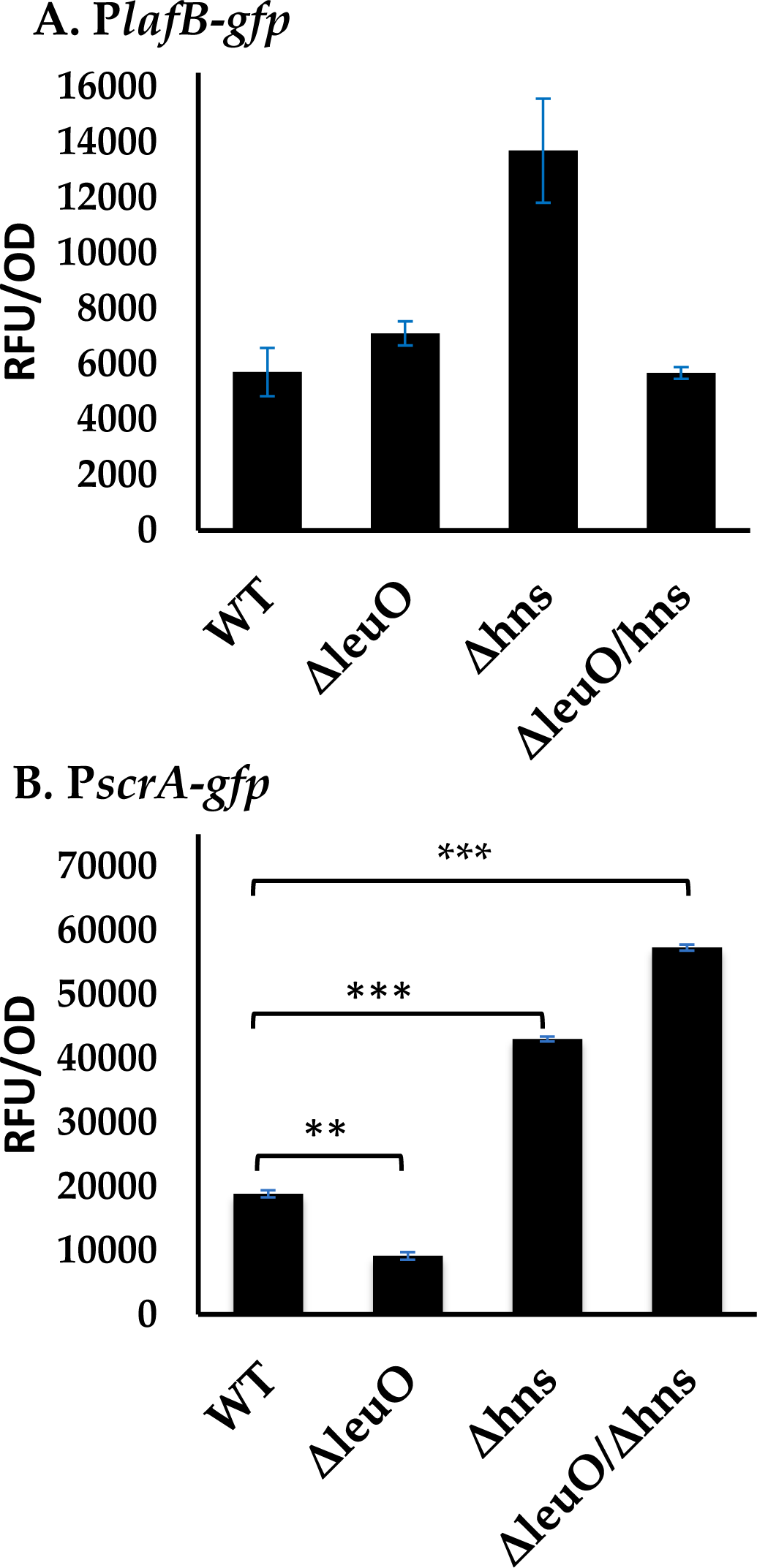
Expression analysis of *lafB* and *scrA*. Reporter assays of **A**. pRUP*lafB*-*gfp* or *B*. pRUP*scrA*-*gfp* in wild-type, Δ*leuO,* Δ*hns, and* Δ*leuO/*Δ*hns* backgrounds. All strains were grown on HI plates 2% NaCl with Tet for 18 h at 30°C. All assays were performed with two biological replicates with three technical replicates. Error bars indicate standard deviation. Unpaired Student’s *t*-test was used to determine the p-value. *p<0.05, **p<0.01, ***p<0.001

### LeuO and HNS control CPS and biofilm production

Enos-Berlage and colleagues described a *V. parahaemolyticus* H-NS deletion mutant as having significant defects in CPS and biofilm production while remaining swarming competent (29). To examine this further, we performed CPS assays with wild type and the three deletion mutant strains (**Fig. 4**). The wild-type strain showed a dense rugose wrinkled colony morphology indicative of CPS production (**Fig. 4**). The Δ*leuO* mutant also showed a rugose crinkly colony morphology but was not as dense as the wild type suggesting less CPS (**Fig. 4**). In contrast, Δ*hns* produced a smooth translucent colony morphology indicating little CPS production (**Fig. 4**). The Δ*leuO*/Δ*hns* strain produced a rough opaque colony morphology. Since CPS is a vital component of biofilm, we investigated how the gene deletions affected biofilm formation at different time points (**Fig. 5**). At all-time points, the Δ*leuO* mutant showed similar biofilm production to wild type, whereas the Δ*hns* mutant showed significant defects in biofilm production compared to wild type. At the 24 h and 48 h time points, the double mutant showed increased biofilm formation compared to wild type and Δ*hns*. These data indicate that H-NS is a positive regulator of CPS and biofilm formation likely by inhibiting *leuO*, a negative regulator (**Fig. 5**).

**Figure 4.**
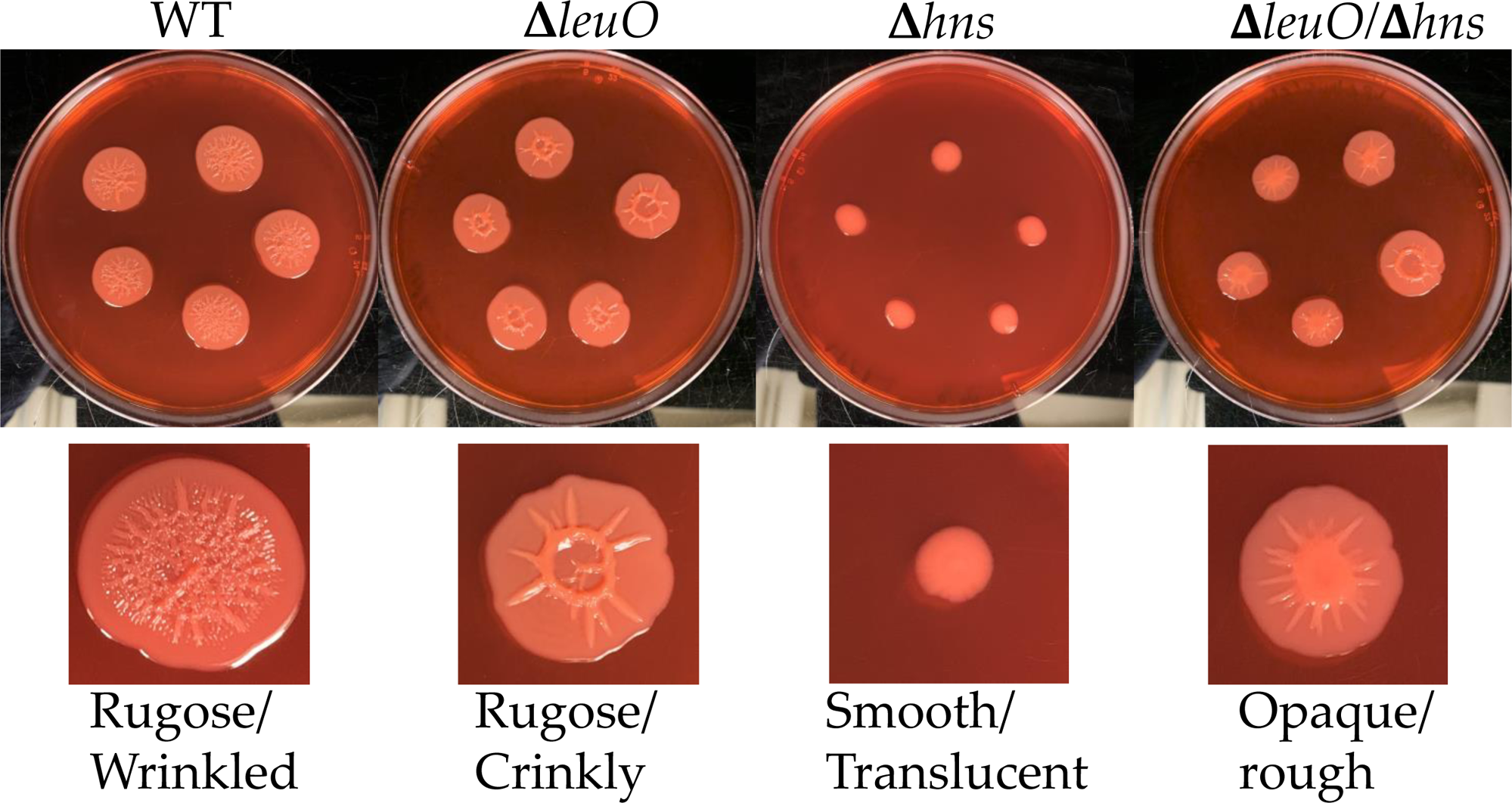
Capsule polysaccharide (CPS) production assays. Wild type (WT), Δ*leuO,* Δ*hns, and* Δ*leuO/*Δ*hns* strains were grown on HI plates supplemented with 0.5% NaCl, CaCl_2_, and Congo Red incubated at 30°C for 48 h. Images are an example of three biological replicates. The structured surface is the result of CPS produced, the more CPS produced, the more wrinkly the colony morphology.

**Figure 5.**
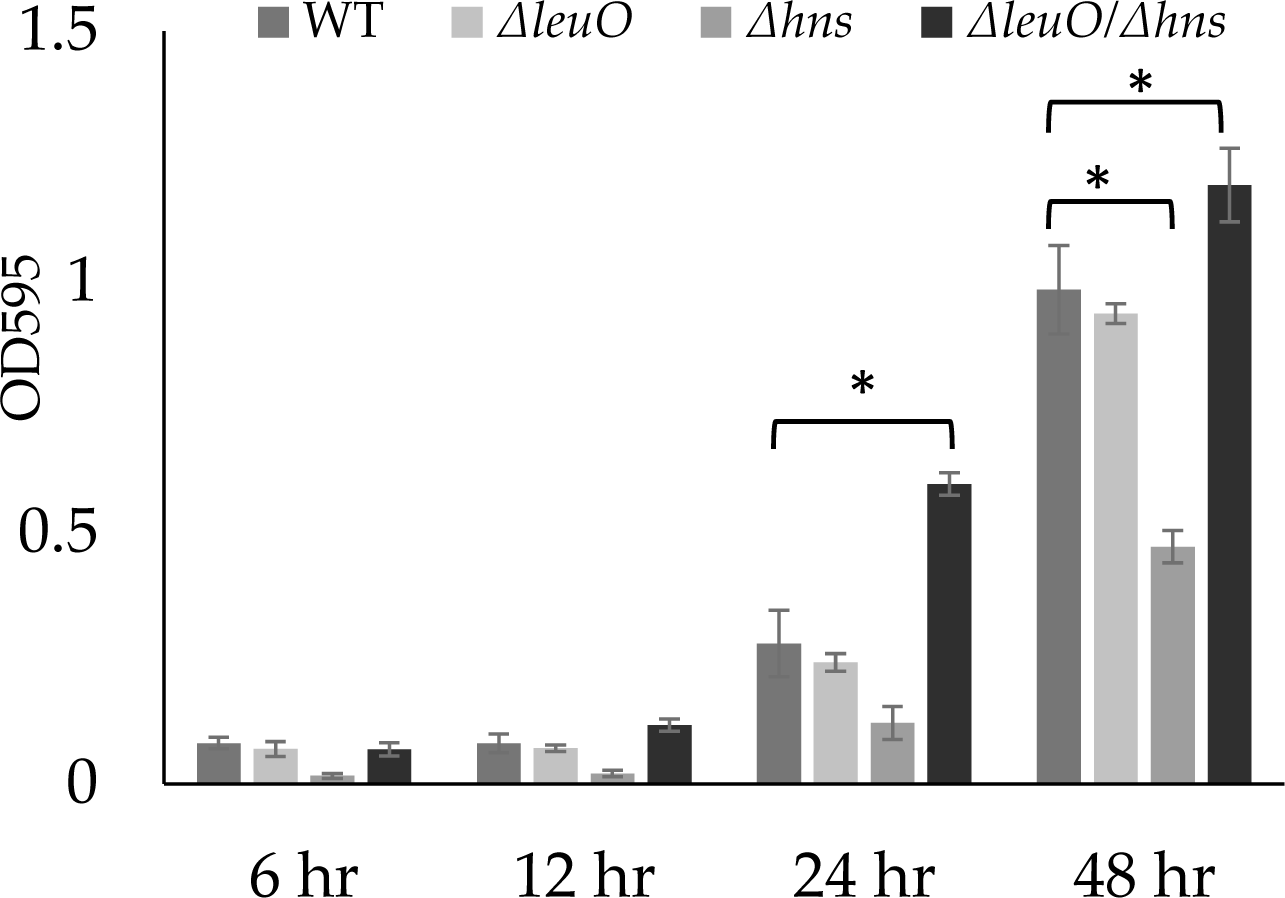
Biofilm assays. Wild type (WT), Δ*leuO,* Δ*hns,* Δ*leuO/*Δ*hns* strains were grown in LB media with 3% NaCl at 30°C with aeration. After each incubation time, biofilms were washed with 1X PBS, stained with 0.1% crystal violet, dissolved in DMSO, and quantified using optical density readings. All assays were performed with two biological replicates with two technical replicates each. Error bars indicate standard deviation. Unpaired t-test was used to determine the p-value. *p<0.05

### LeuO and H-NS regulate *cpsA* expression

To determine how H-NS and LeuO control CPS and biofilm production, the promoter region of the *cps* operon was cloned into pRUP164 and P*cpsA*-*gfp* fluorescence measured in wild type, Δ*leuO,* Δ*hns,* and Δ*leuO/*Δ*hns*. Reporter assays were performed on cells grown on HI media plates with 0.5% NaCl Tet for 48 h at 30°C. P*cpsA*-*gfp* in the Δ*leuO* strain showed significantly increased fluorescence whereas in Δ*hns* fluorescence was significantly reduced compared to wild type (**Fig. 6A**). In the Δ*leuO*/Δ*hns* strain fluorescence was significantly increased (**Fig. 6A**). These data show that H-NS is a positive regulator of the *cps* operon whereas LeuO is a negative regulator. At the *cpsA* locus, LeuO is epistatic to H-NS as deletion of both genes resulted in higher fluorescence than wild-type.

**Figure 6.**
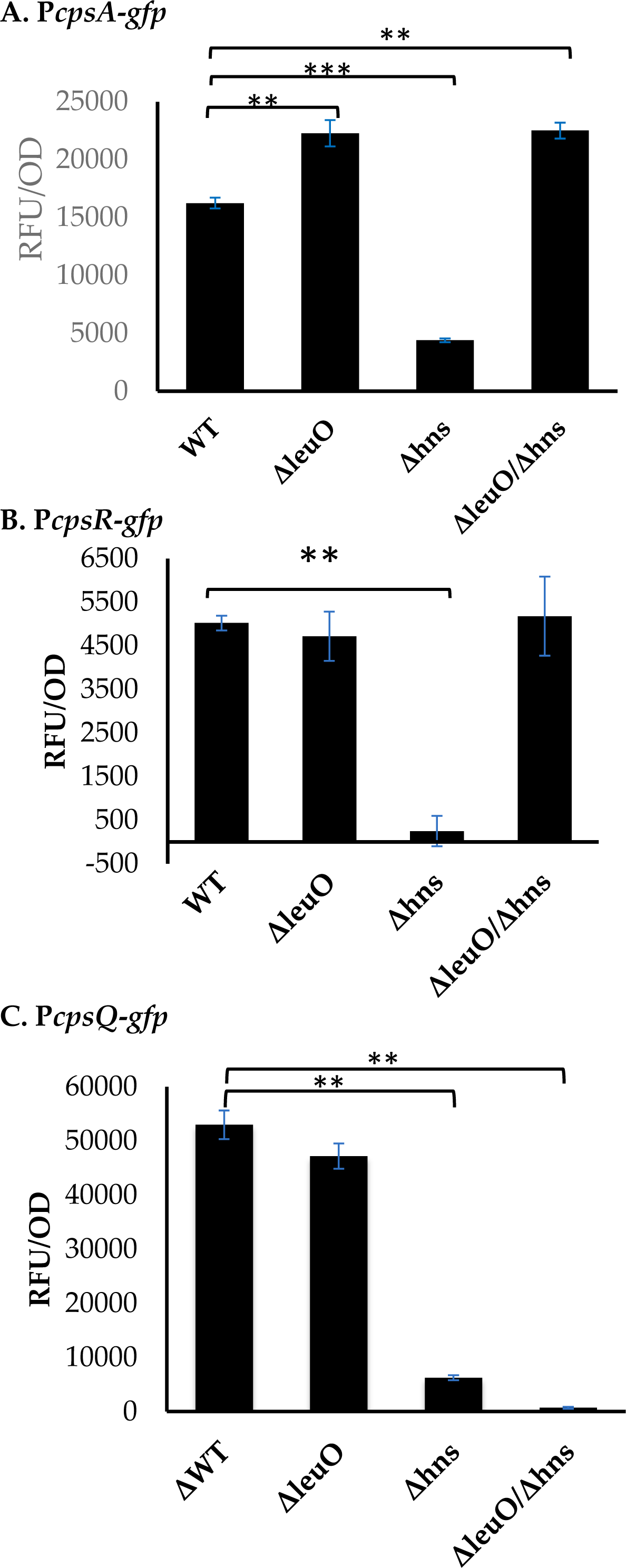
Expression analysis of *cpsA, cpsR* and *cpsQ*. Reporter assays with pRUP*cpsA*-*gfp,* pRUP*cpsR*-*gfp,* or pRUP*cpsQ*-*gfp* in wild type (WT), Δ*leuO,* Δ*hns,* Δ*leuO/*Δ*hns* backgrounds. All strains were grown on HI 0.5% NaCl plates supplemented with Tet and at 30°C for 18 h. All assays were performed with two biological replicates with three technical replicates. Error bars indicate standard deviation. Unpaired t-test was used to determine the p-value. **p<0.01, ***p<0.001

Next, we examined two regulators of the *cps* operon, CpsR encoded by *cpsR*, is a positive regulator of *cpsQ*, which encodes the c-di-GMP dependent regulator CpsQ, a direct positive regulator of the *cps* operon (14, 16). The promoter regions of *cpsR* and *cpsQ* were each cloned upstream of the promoterless *gfp* gene in pRUP164 and transformed into wild-type and the three deletion mutants (**Fig. 6B and 6C**). P*cpsR*-*gfp* fluorescence in Δ*leuO* background was similar to wild-type whereas in Δ*hns* significantly decreased fluorescence was observed. In the double-mutant, fluorescence reverted to wild-type and Δ*leuO* levels (**Fig. 6B**). This indicates an epistatic relationship between H-NS and LeuO at the *cpsR* locus. In P*cpsQ*-*gfp* assays, fluorescence in Δ*leuO* was similar to wild-type, but in both Δ*hns* and Δ*leuO*/Δ*hns* backgrounds significantly decreased fluorescence was observed (**Fig. 6C**). The data show H-NS is a positive regulator, but LeuO does not play a role at the *cpsQ* locus.

## DISCUSSION

Here, we investigated the role of H-NS and LeuO in controlling the decision between swarming motility and sessile biofilm formation, two mutually exclusive group behaviors present in *V. parahaemolyticus*. The data showed that both regulators played a role in these phenotypes with H-NS a negative regulator of swarming motility and a positive regulator of CPS and biofilm formation (**Fig. 7**). The data suggest that H-NS impacted LeuO, in both swarming and CPS and biofilm formation. The single mutants suggested that H-NS had a more significant role compared to LeuO, as the Δ*hns* mutant showed more apparent phenotypes in all assays whereas the Δ*leuO* mutant showed much more subtle differences compared to wild type. The analysis of the double mutant uncovered the significance of LeuO and showed it was epistatic to H-NS in the control of both phenotypes playing a role downstream of H-NS. In most of our experiments (phenotypes and expression), an epistatic affect was present, and in most cases, the Δ*hns* mutant phenotypes were lost in the double mutant. The clear exception was the *cpsQ* locus, which was regulated by H-NS, but not LeuO.

**Figure 7.**
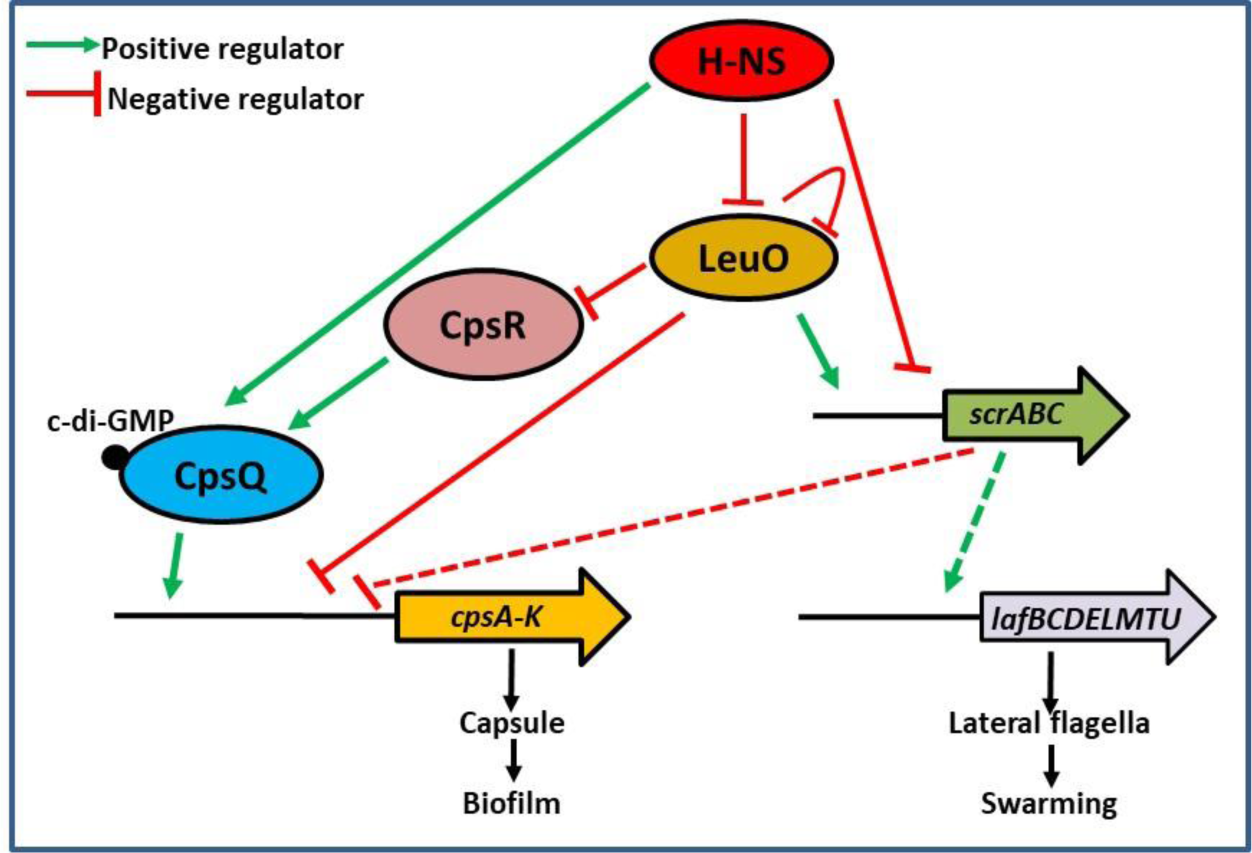
Proposed model of LeuO and H-NS oppositely regulating swarming motility and capsule production in *V. parahaemolyticus.* LeuO is a negative auto-regulator and is also negatively regulated by H-NS. LeuO positively regulates the *scrABC* operon, but is a negative regulator *cpsA* and *cpsR* expression. H-NS acts as a negative regulator of swarming motility and a positive regulator of CPS production, in part by controlling LeuO. H-NS positively regulates *cpsQ* expression independent of LeuO.

H-NS and LeuO have been well studied for their role as global gene regulators and their antagonistic relationship at many loci in *E. coli*, *Salmonella spp.,* and *V. cholerae* (37, 38, 40–43). In *Vibrio* species, H-NS has been studied for its involvement in virulence and bioluminescence, but much less is known about its interactions with LeuO. In *V. vulnificus*, a human pathogen, the RTX toxin, an important virulence factor, is repressed by H-NS and is an inhibitor of cytotoxicity by repressing hemolysin production (50). H-NS is also known to silence LeuO, the master regulator of the cyclic (PhePro)-dependent signal pathway in *V. vulnificus* (51). In *V. harveyi*, H-NS represses the *luxCDABE* operon responsible for bioluminescence in a cell concentration dependent manner (52). In *V. cholerae*, H-NS and LeuO were shown to work together to repress the *vieSAB* loci, which encode a phosphodiesterase involved in biofilm production and motility (42, 43). It is of interest to note that although *V. cholerae* contains many of the same regulators of CPS and biofilm as *V. parahaemolyticus*, their effects are the opposite between the two species (25–27). In *V. cholerae,* the *vpsA-K, vpsU, and vpsL-Q* operons, which produce CPS, are repressed by H-NS under low c-di-GMP conditions (41, 42, 53). HapR the quorum sensing master regulator negatively regulates *vpsT* and *vpsR*, two direct positive regulators of *vps* production (54). H-NS is also a negative regulator of *vps* by negatively regulating *vpsT,* and CRP, the cAMP regulator protein, is a negative regulator of *vps* expression (25–27, 55–57). In *V. parahaemolyticus*, studies have shown that H-NS is a positive regulator of biofilm and CPS, and a negative regulator of swarming motility (29, 44, 45). The present study shows that H-NS regulation of these phenotypes involves LeuO and their regulation of swarming genes *laf* and *scrABC* and biofilm genes *cpsA-K* and their regulator *cpsR*. In *V. parahaemolyticus*, it was shown that OpaR (the HapR homolog) is a positive regulator of CPS production and a negative regulator of swarming motility due to the positive effects of OpaR on c-di-GMP levels and *cps* expression (12, 28–30, 58–63) (64). In a Δ*opaR* mutant, the expression of *cpsA*, *cpsR,* and *cpsQ* is downregulated in *V. parahaemolyticus* indicating OpaR is a positive regulator of CPS and biofilm formation (14, 60). In support of this positive role in CPS and biofilm, in the Δ*opaR,* both the *laf* and *scrABC* operons were highly expressed and cells produce hyper swarming cells showing OpaR is a negative regulator of swarming motility (10, 30, 61, 63). A recent study showed that a *crp* deletion mutant, which encodes CRP, had a defect in CPS and biofilm formation indicating a positive role in this phenotype (65).

In conclusion, our study has demonstrated the importance of H-NS and LeuO in the control of the decision between the surface attachment behaviors swarming and biofilm formation. Our work suggests an epistatic relationship between LeuO and H-NS with H-NS in general playing a silencing role to oppositely control swarming and biofilm formation. Given the central role of OpaR and CRP in biofilm formation it will be of interest to learn what role H-NS plays in the regulation of these global regulators.

## METHODS

### Bacterial Strains, Media, and Growth Conditions

A list of all strains used in this study is found in Table 1. *Vibrio parahaemolyticus* RIMD2210633, a streptomycin-resistant clinical strain, was used as wild type (WT) in all experiments (66, 67). *Escherichia coli* DH5∝ and *E. coli β*2155 *λ*pir (diaminopimelic acid auxotroph) strains were used for cloning experiments. LB media with 3% NaCl (LBS) was used when growing *V. parahaemolyticus* strains unless stated otherwise. LB media was used when growing *E. coli* strains and supplemented with 0.12 mM diaminopimelic acid (DAP) when growing *E. coli β*2155 *λ*pir. Antibiotics streptomycin (Sm, 200 μg/ml), chloramphenicol (Cm, 12.5 μg/ml), and tetracycline (1 μg/ml) were used.

**Table 1.** Bacterial strains used in this study.

| Strain or plasmid | Genotype or description | Reference(s) or source |
| --- | --- | --- |
| ***Vibrio parahaemolyticus*** |  |  |
| RIMD2210633 | O3:K6 clinical isolate, Str<sup>r</sup> | (66) |
| $\Delta leuO$ | RIMD2210633 $\Delta leuO$ (VP0350), Str<sup>r</sup> | (47) |
| $\Delta hns$ | RIMD2210633 $\Delta hns$ (VP1133), Str<sup>r</sup> | (48) |
| $\Delta leuO/\Delta hns$ | RIMD2210633 $\Delta leuO \Delta hns$ (VP0350/VP1133), Str<sup>r</sup> | (48) |
| ***Escherichia coli*** |  |  |
| DH5α λpir | $\Delta lac$ pir | ThermoFisher Scientific |
| B2155 λpir | $\Delta dapA::erm$ pir for bacterial conjugation | (68) |
| **Plasmids** |  |  |
| pRU1064 | promoterless-*gfp* UV Amp<sup>r</sup>, Tet<sup>r</sup>; IncP ori | (69) |
| pRUP$cpsA$-*gfp* | pRU1064 with *cpsA* promoter Amp<sup>r</sup>, Tet<sup>r</sup> | This study |
| pRUP$cpsR$-*gfp* | pRU1064 with *cpsR* promoter Amp<sup>r</sup>, Tet<sup>r</sup> | This study |
| pRUP$cpsQ$-*gfp* | pRU1064 with *cpsQ* promoter Amp<sup>r</sup>, Tet<sup>r</sup> | This study |
| pRUP$scrA$-*gfp* | pRU1064 with *scrA* promoter, Amp<sup>r</sup>, Tet<sup>r</sup> | (63) |
| pRUP$lafB$-*gfp* | pRU1064 with *lafB* promoter, Amp<sup>r</sup>, Tet<sup>r</sup> | (63) |
| pRUP$leuO$-*gfp* | pRU1064 with *leuO* promoter Amp<sup>r</sup>, Tet<sup>r</sup> | (48) |

### Swimming and swarming motility assay

*V. parahaemolyticus* strains were grown on LBS plates overnight and colonies used to spot inoculate semisolid LB 2% NaCl 0.3% agar plates incubated at 37°C for 24 h. The swimming diameters for each strain was measured and plate were imaged. Heart Infusion (HI) media plates were prepared with 2% NaCl and 1.5% agar. Single colonies were used to spot inoculate each HI plate at four positions and incubated at 30°C for 48 h. Plates were imaged, and swarming diameter was measured. The assays were performed using at three biological replicates and three technical replicates for each strain.

### Capsular polysaccharide (CPS) and biofilm assays

To observe CPS, HI media plates were prepared with 0.5% NaCl, 0.04% CaCl_2_, and 0.25% Congo red dye. Single colonies from overnight LBS plates were used to spot inoculate each HI plate at five positions and were incubated at 30°C for 48 h, after which plates were imaged. Biofilm assays were performed using 8-well plates strips containing 200 µL LBS media incubated with shaking at 30°C for 6 h, 12 h, 24 h, and 48 h. At the end of each incubation time point, liquid media was discarded, and the wells were washed with 1x PBS twice. 0.1% crystal violet was then added to each well and the wells were incubated at room temperature for 30 mins. After incubation, crystal violet was discarded, and wells were washed again with 1x PBS thrice. Wells were then imaged to visualize the stained biofilms, followed by solubilization of the biofilms in 220 µL DMSO and quantified via OD_595_ absorbance readings. Biofilm assays were performed with at least two biological replicates with two technical replications.

### Transcription reporter vector construction

pRU1064 plasmid with a promoterless *gfp* gene and a tetracycline resistance gene was used as a reporter vector. Reporter vectors were constructed by cloning the promoter regions of *leuO*, *lafB, scrABC, cpsA-K, cpsR,* and *cpsQ,* into the SpeI site upstream of the *gfp* gene via Gibson Assembly. Gibson primers were designed for the promoter regions of interest and were used to amplify the regions of interest using high-fidelity Phusion polymerase in PCR reactions. Overnight cultures of *E. coli* pRU1064 were used to isolate and purify pRU1064, which was then digested with SpeI. The digested plasmid and amplified promoter region PCR products were incubated together with 2x Hifi buffer at 50°C for 30 mins to create to create pRUP*_leuO_*-*gfp*, pRUP*_cpsA_*-*gfp*, pRUP*_cpsR_*-*gfp*, pRUP*_cpsQ_*-*gfp,* pRUP*_lafB_*-*gfp,* and pRUP*_scrA_*-*gfp*. Each plasmid was transformed into *E. coli* DH5*ɑ* competent cells by heat shock. Transformed cells were grown on LB media with tetracycline and screened for the presence of the plasmid. Positive colonies were used to set up overnight cultures, which were used to extract the purified plasmid. The purified plasmid was used to transform *E. coli β*2155, which was plated on LB media with chloramphenicol and DAP and screened in the same manner. Conjugations were set up with *E. coli β*2155 and *V. parahaemolyticus* strains on LB media with tetracycline. Colonies were screened for the presence of the required plasmid with PCR using primers for the multiple cloning site followed by gel-electrophoresis.

**Table 2.**
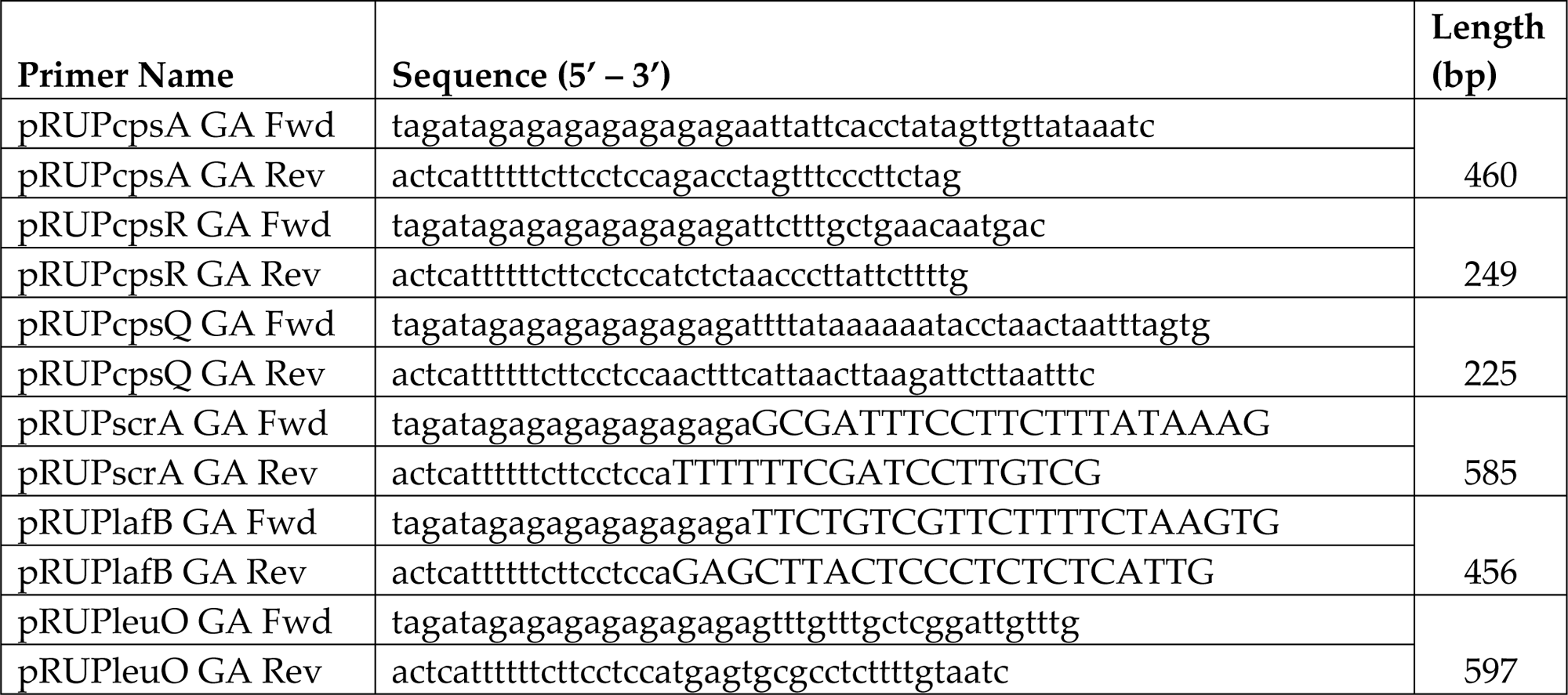
Reporter primer pairs used in this study.

### Transcription reporter assays

*V. parahaemolyticus* pRUP*_lafB_*-*gfp,* and pRUP*_scrA_*-*gfp* were spot inoculated on HI 2% NaCl swarming agar plates and incubated for 18 h at 30°C. *V. parahaemolyticus* pRUP*cpsA*, pRUP*cpsQ*, or pRUP*cpsR* were spot inoculated on HI media 0.5% NaCl plates and incubated for 18 h at 30°C. Colonies were scraped off the plate, washed and suspended in 1x PBS. OD_600_ was measured after resuspension and was adjusted to final OD of 0.5. GFP fluorescence were measured in these cells with excitation at 385 nm and emission at 509 nm in black, clear-bottom 96-well plates on a Spark microplate reader with Magellan software (Tecan Systems Inc.). Specific fluorescence was calculated by dividing relative fluorescence for each sample by its OD_595_ reading. Two biological replicates were performed per sample in triplicate. Student’s *t*-test was used to calculate significance.

